# Criticality as a determinant of integrated information Φ in human brain networks

**DOI:** 10.1101/720151

**Authors:** Hyoungkyu Kim, Anthony G. Hudetz, George A. Mashour, UnCheol Lee

## Abstract

Integrated information theory (IIT) postulates that consciousness arises from the cause-effect structure of a system but the optimal network conditions for this structure have not been elucidated. In the study, we test the hypothesis that network criticality, a dynamically balanced state between a large variation of functional network configurations and a large constraint of structural network configurations, is a necessary condition for the emergence of a cause-effect structure that results in a large Φ, a surrogate of integrated information. We also hypothesized that if the brain deviates from criticality, the cause-effect structure is obscured and Φ diminishes. We tested these hypotheses with a large-scale brain network model and high-density electroencephalography (EEG) acquired during various levels of human consciousness during general anesthesia. In the modeling study, maximal criticality coincided with maximal Φ. The constraint of the structural network on the functional network is maximized in the maximal criticality. The EEG study demonstrated an explicit relationship between Φ, criticality, and level of consciousness. Functional brain network significantly correlated with structural brain network only in conscious states. The results support the hypothesis that network criticality maximizes Φ.

## Introduction

Integrated information theory (IIT) proposes that consciousness equates with integrated information in a system and the integrated information is maximized when integration and differentiation of the systems’ components are balanced. IIT proposes algorithmic methods to identify the differentiated parts of a system and quantify the integrated information across the parts [1–6]. Φ is a measure of complexity of the cause-effect structure of the minimum information partition among all possible partitions. However, in a dynamic system such as the brain, the optimal conditions under which the cause-effect structure—that is, the basis of integrated information— arises has not been elucidated. In this study, we hypothesized that network criticality, a balanced state between a large variation of functional network configurations and a large constraint of structural network configurations, may be the basis of the high Φ in conscious brains.

Criticality was originally introduced for studying phase transition in physics, which was simply defined as a balanced state between order and disorder in the activities of the elements that make up a system [7,8]. This property has been observed broadly in physical and non-physical systems and has been suggested as an optimal state for information storage, transmission, and integration with high susceptibility to external stimuli [9–13]. In particular, several computational modeling and empirical studies suggest that the brain dynamics associated with consciousness reside near a critical state [14–19]. Furthermore, recent studies have attempted not only to identify whether the conscious brain resides near the critical state but also to understand systematically how various types of brain perturbations (sleep, anesthesia, and traumatic injuries) lead to a deviation from criticality [9,20–23]. Such approaches introduce the criticality hypothesis as a theoretical framework to study pharmacological and pathological states of unconsciousness, such as anesthesia and coma. Regarding the characteristics of a critical state such as highly informative, highly susceptible, highly efficient, and highly integrative, criticality shares many commonalities with the brain state that IIT proposes as conducive to consciousness. However, the relationship between criticality, Φ, and human consciousness has not been demonstrated explicitly.

To identify a relationship between criticality and Φ, we analyzed computationally a large-scale human brain network model adjusting criticality with a control parameter. The criticality was defined using the pair correlation function (PCF), a surrogate measure of susceptibility [24]. The relative change of Φ across states was estimated with 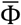, a surrogate measure of Φ that we developed for high-density EEG. Empirically, we modulated the level of human consciousness in a stepwise manner with an anesthetic and calculated both 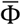 and PCF from continuous EEG data. From the modeling study and EEG analysis, we were able to quantitatively study the relationship between Φ, criticality, and consciousness, and suggested network criticality as a necessary condition for a high Φ in human brain networks.

## Methods

### Ethics Statement

This study was conducted at the University of Michigan Medical School and approved by the Institutional Review Board (HUM00061087); after careful discussion, written informed consent was obtained from all participants.

### Human brain network model

Many recent studies have successfully applied Kuramoto/Stuart-Landau models to the brain in order to understand the organizational principles of multiscale brain function, surrogates of information flow, and complex dynamics at the whole brain network level [25–27]. Similarly, we hypothesized that the application of simple oscillatory models, which can modulate the criticality with a control parameter, to an anatomically informed brain network structure could inform the relationship between brain network criticality and integrated information.

We used a large-scale brain network model that implements a coupled simple oscillator model on the scaffold of an anatomically informed human brain network structure. The human brain network consists of 78 parcels of the cerebral cortex constructed from diffusion tensor imaging (DTI) of 80 young adults [28].

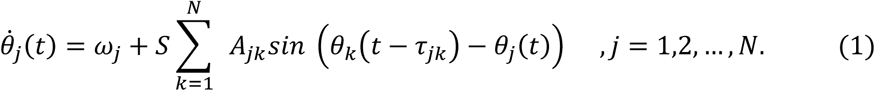

Here *S* is the coupling strength between oscillators and A_*jk*_ denotes the anatomical connections between oscillator *j* and *k*, yielding 1 if a connection exists and 0 otherwise. *τ*_*jk*_ is the time delay between node *j* and *k. θ*_*j*_*(t)* is the phase of oscillator *j* at time *t. ω*_*j*_ is the intrinsic frequency of oscillator *j*.

### Simulation procedures

All parameters for the models were set to simulate alpha oscillations in the brain. Alpha oscillation models have successfully explained empirically observed brain network behaviors such as functional connectivity, traveling waves, and network state transitions based on electroencephalography (EEG) and magnetoencephalography (MEG) [25,26,29–37]. Thus, alpha oscillations were analyzed to understand the behaviors of Φ and the criticality at the brain network level. The natural frequencies of the oscillators in our simulation were given as a Gaussian distribution around 10 Hz with a standard deviation of 0.5 Hz. Time delay was set proportional to the physical distances between nodes with a propagation speed of 8.6m/s [38]. The coupling strength between the oscillators was continuously changed from 0 to 18. In each parameter set, 100 configurations were simulated and the results were averaged over all configurations.

### Experimental procedures

We conducted a secondary analysis of high-density EEG data from a study of sevoflurane-induced unconsciousness in humans; the details of the experiment can be found in the supplementary text of this article and the previously published Blain-Moraes et al [39]. The current study tested different hypotheses and, unlike the original study, included computational analyses of brain network models.

### Sevoflurane dataset

64-channel EEG were recorded from seven healthy volunteers as sevoflurane concentrations in high-flow oxygen (8 L/min) were gradually increased from 0.4% to 0.6% to 0.8% (the average range at which unconsciousness was induced) or beyond, then decreased from 0.8% to 0.6% to 0.4%. The EEG was recorded with eyes closed. The loss and recovery of consciousness were defined as the loss and recovery of response to the verbal command ‘squeeze your left [or right] hand twice,’ on a recorded loop every 30 seconds, with right/left hand commands randomized.

Average reference was used for re-referencing and the windowed sinc-FIR filter (in the MATLAB toolbox from EEGLAB) was used to avoid a possible shifting of the signal. We analyzed 12-second-long EEG epochs with 1-minute-long moving windows.

### Criticality

Criticality, a boundary state between order and disorder, has long been proposed to play an important role in neural dynamics and brain function. Empirical evidence supports the hypothesis that the brain operates at or near the critical point, not only at the neuronal network level [22,40,41], but also at the large-scale or global network level [9,12,42–44]. Until now, most studies have focused on scale-free behavior, showing power law distribution of empirically observed variables. It has also been recently proposed that high correlation between functional and structural brain networks [14,23,25,26,45] and a large pair correlation function (PCF) [24,46] is evidence of criticality. In both the brain network model and empirical EEG data, we estimated criticality with PCF, which is the variance of global phase synchronization and is equivalent to susceptibility in statistical physics

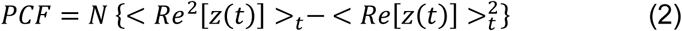

where Re[*z(t)*] is the real part of the *z(t)* in Eq. (3).

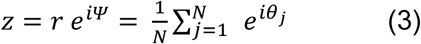

where Ψ is the order parameter phase. The absolute value *r* = |z| represents the degree of synchronization. The *r* is equal to zero when the phases of nodes are uniformly distributed and one when all the nodes have the same phase.

### Calculation of 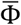

IIT defines integrated information (Φ) as the effective information (*φ*) of the minimum information partition (MIP) in a system [1–3,5,47]. The MIP is also defined as the partition having minimum effective information among all possible partitions.

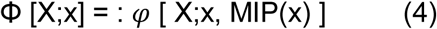

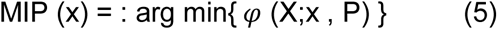

where X is the system, x is a state, and P is a partition P={*M*^1^, …, *M*^*r*^}.

Identifying the MIP requires searching all possible partitions and comparing their effective information φ. This is the most time-consuming and computationally demanding process in the application to high-density EEG. Furthermore, considering the fact that EEG data recorded during anesthetic state transitions are non-Gaussian and continuous time series, we used the 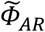 as a measure of integrated information [48]. The computed 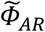 values from original signals were compared with the 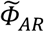 values of surrogate data sets, and non-significant 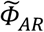 values were set as zero.

Although many improvements have been made in the algorithms of Φ over the last decade, the computation time is still unrealistic because of the need to search an enormous number of partitions to identify the MIP. In our previous study, we proposed a surrogate measure 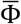 to circumvent the explosive computational time of Φ for high-density EEGs. 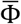 estimates the relative change of Φ across states by considering the average feature of many small sample units rather than trying to identify the MIP and its effective information for all EEG channels. A sample unit consists of a small number of EEG channels randomly selected from 64 channels and the total number of sample units is taken as large enough to represent the behavior of the entire high-density EEG montage. In this study, we limited our interest only to the relative changes of Φ values across states, rather than attempting to calculate the exact Φ value for each brain state, which would be impossible to measure using the superficial and spatially imprecise brain activity recorded by EEGs.

For each sample unit, we were able to calculate the MIP and its effective information—that is, the Φ of the sample unit. For instance, in this study, we selected 8 random EEG channels for a sample unit and acquired 200 sample units that were randomly selected from the baseline states. The same 200 sample units determined in the baseline were then compared across EEG windows to investigate the increase or decrease of Φ values. Since the number of possible bipartitions of 8 channels is 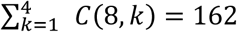, where *C* stands for the combination of *k* unique elements chosen from eight possible elements, calculating Φ for all EEG windows of seven subjects during state transitions is possible within a relatively short computational time.

The average 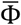 is defined as follows.

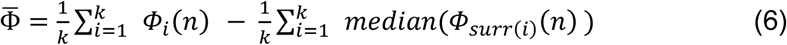

where *n* is the number of EEG channels for a sample unit and *k* is the number of sample units of the *n* EEG channels. *Φ*_*i*_(*n*) measures the effective information of MIP for the *n* EEG channels, by definition, the integrated information of the sample unit. Here, we chose n=8 and k=200 following our previous study [49]. *Φ*_*surr*_(*n*) is the spurious *Φ*_*i*_(*n*) estimated from randomly shuffled EEG data sets. Subtracting the randomness, 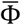 reflects the average integrated information of 200 sample units taken from the high-density EEG data that exceeds the spurious information integrated from the surrogate data. Since 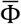 estimates the relative change of Φ, it is appropriate for our purposes to detect the maximum Φ to compare with the maximum criticality.

### Functional connectivity

Phase Lag Index (*PLI*), a measure of phase locking between two signals, was used to define the functional connectivity in the EEG network [50]. We chose a Hilbert transform to extract the instantaneous phase of the EEG from each channel and calculate the phase difference *Δθ*_*ij*_(*t*) between channels *i* and *j*, where *Δθ*_*ij*_(*t*) = *θ*_*i*_(*t*) – *θ*_*j*_(*t*), *t = 1,2,…,n*, and *n* is the number of samples within one epoch. *PLI*_*ij*_ between two nodes *i* and *j* is then calculated using equation (7):

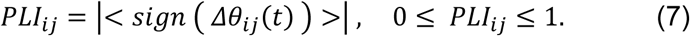

Here, the sign() function yields: 1 if *Δθ*_*ij*_(*t*) > 0; 0 if *Δθ*_*ij*_(*t*) = 0; and −1 if *Δθ*_*ij*_(*t*) < 0. The mean < > is taken over all *t=1,2,…,n*. If the instantaneous phase of one signal is consistently ahead of the other signal, the phases are considered locked and *PLI*_*ij*_ ≈ 1. However, if the signals randomly alternate between a phase lead and phase lag relationship, there is no phase locking and *PLI*_*ij*_ ≈ 0.

### Surrogate data

To control for spurious connectivity of EEG, 20 surrogate data sets were generated with a random shuffling method, in which a time point is randomly chosen in each EEG channel; the EEG epochs are then shuffled before and after the time point. The shuffled data have the same amplitude distribution and power spectrum of the original EEG, but there are disruptions of the original connectivity between two EEG signals.

### Network construction

We expected that different EEG frequency bands and different states would have different levels of spurious connectivity [51]. Thus, after subtracting the median *PLI* of 20 surrogate data sets, if the remaining *PLI* was larger than 0.1 then the connectivity of two EEG signals was set as 1; otherwise, it was set as 0. The threshold (0.1) was chosen to avoid isolated nodes in the EEG network in the baseline states. The node degree of an EEG channel was defined as the number of functional links in the network.

### EEG simulation

To test if the *PLI* network of EEG in conscious states is similar to the structural brain network in a critical state, we simulated 78 source signals in the structural brain network in a critical state and projected it into the 64 sensor signals on the scalp. We could then directly compare the *PLI* networks of EEG and the *PLI* network of the simulated EEG (; the 64 sensor signals on the scalp). The signals generated from the Kuramoto model and structural brain network represent source activities of the brain, which are under the surface of where EEG is measured. In reality, the electrical potentials generated by the neural activity in the brain conduct outwards through brain tissue and the skull and finally appear at the scalp surface where the EEG signal is measured. In order to compare experimental EEG and the model signals, we generated surface level signals from the simulated source signals. We used three concentric spherical head models; the three layers consist of the brain, skull, and scalp. The conductivity of the three layers was set to be 0.33, 0.0042, and 0.33 S/m, respectively [52]. The source activity was represented as a dipole moment. The coordinate of the dipole moment in the brain was determined by the region’s standard coordinates and the orientation of the dipole moment was randomly assigned. The forward model simulation was conducted by using the Field Trip Toolboox [53].

### Statistical Analysis

We performed one-way ANOVA (“anova1.m”, MATLAB toolbox) with Tukey-Kramer correction (“multcompare.m” with alpha = 0.05 and ctype = “tukey-kramer” in MATLAB) for the comparison of the PCF and 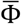 among different states. The statistical tests were carried out for the modeling and empirical analysis separately. The adjusted P-values of 0.05 or lower (*P < 0.05, **P < 0.01, and ***P < 0.001) were considered to be statistically significant (S1 Table and S2 Table).

## Results

### Model Study: Correlation between Criticality and 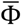

We simulated various brain network dynamics by adjusting the control parameter (i.e., coupling strength). First, we determined the critical state of the model by identifying the coupling strength that yielded a maximum criticality. The criticality of the system was defined by PCF, which reflects the susceptibility of the brain network dynamics to perturbation. As the coupling strength of the network was increased, the PCF reached a maximum at an intermediate coupling strength, while the order parameter increased monotonically (dotted line in Figure 2A). In Figure 2A, we illustrate the intermediate coupling strength (blue region) as the critical state of this brain network model and selected an incoherent (green region) and highly synchronized (red region) states of lower PCFs as a supercritical and subcritical state, respectively. The 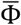 value was also maximized at a point between sub- and supercritical states, at a similar coupling strength that corresponds to the maximum PCF.

**Figure 1.**
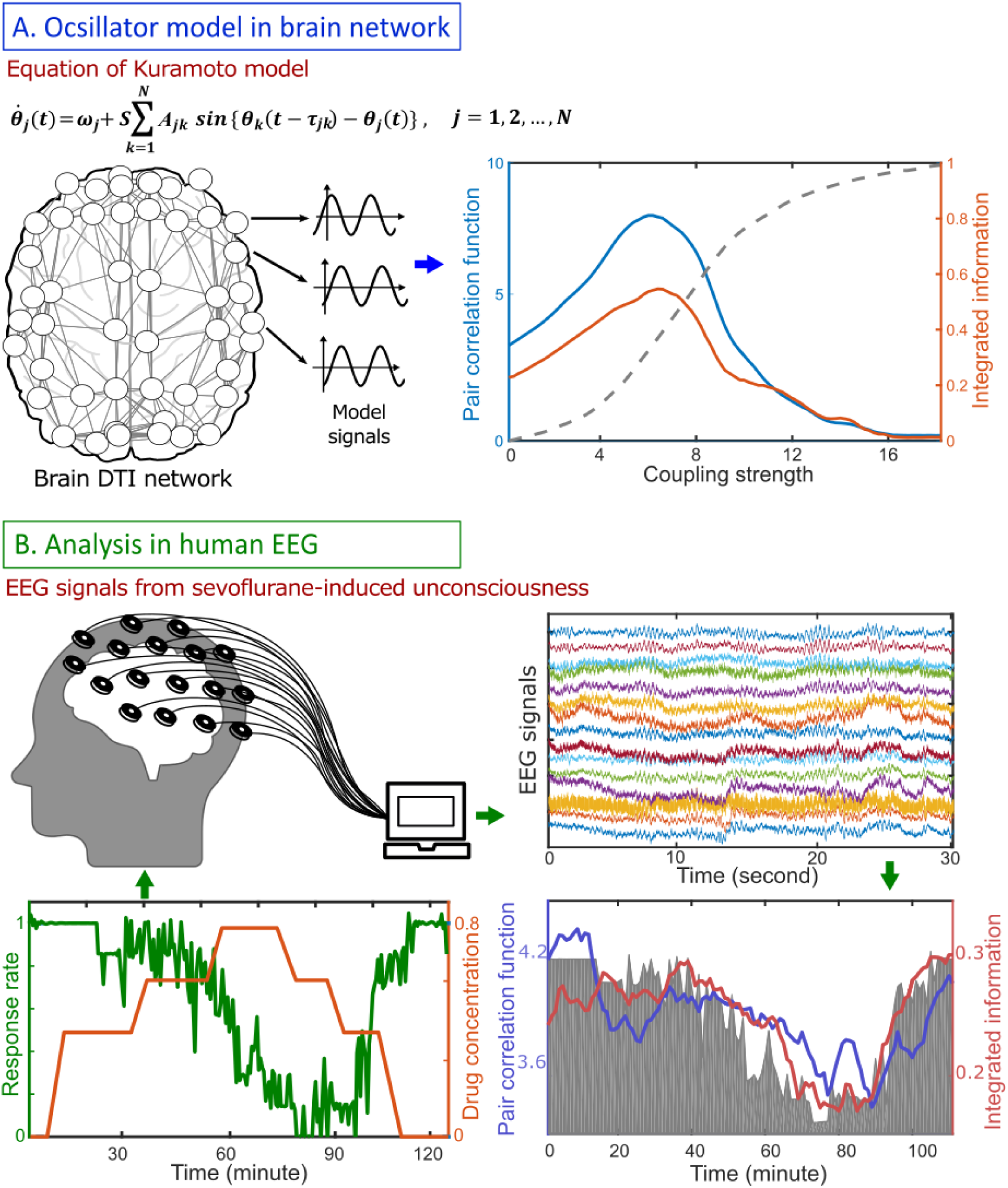
Schematic flow diagram for studying the relationship between criticality (defined as pair correlation function, PCF), integrated information (Φ) and human consciousness. (A) Model study: a simple coupled oscillator model (Kuramoto model) was implemented on an anatomically informed human brain network structure constructed from diffusion tensor imaging (DTI). The Φ was calculated while modulating the level of criticality with a control parameter (coupling strength). The level of criticality was defined using PCF, a susceptibility measure, of the simulated brain activity. (B) Empirical study: 64-channel EEG data derived from seven healthy volunteers were recorded while gradually increasing sevoflurane concentration from 0.4% to 0.6% to 0.8%, then decreasing it from 0.8% to 0.6% to 0.4%. The changes of PCF and Φ were compared with the response rate to verbal commands.

**Figure 2.**
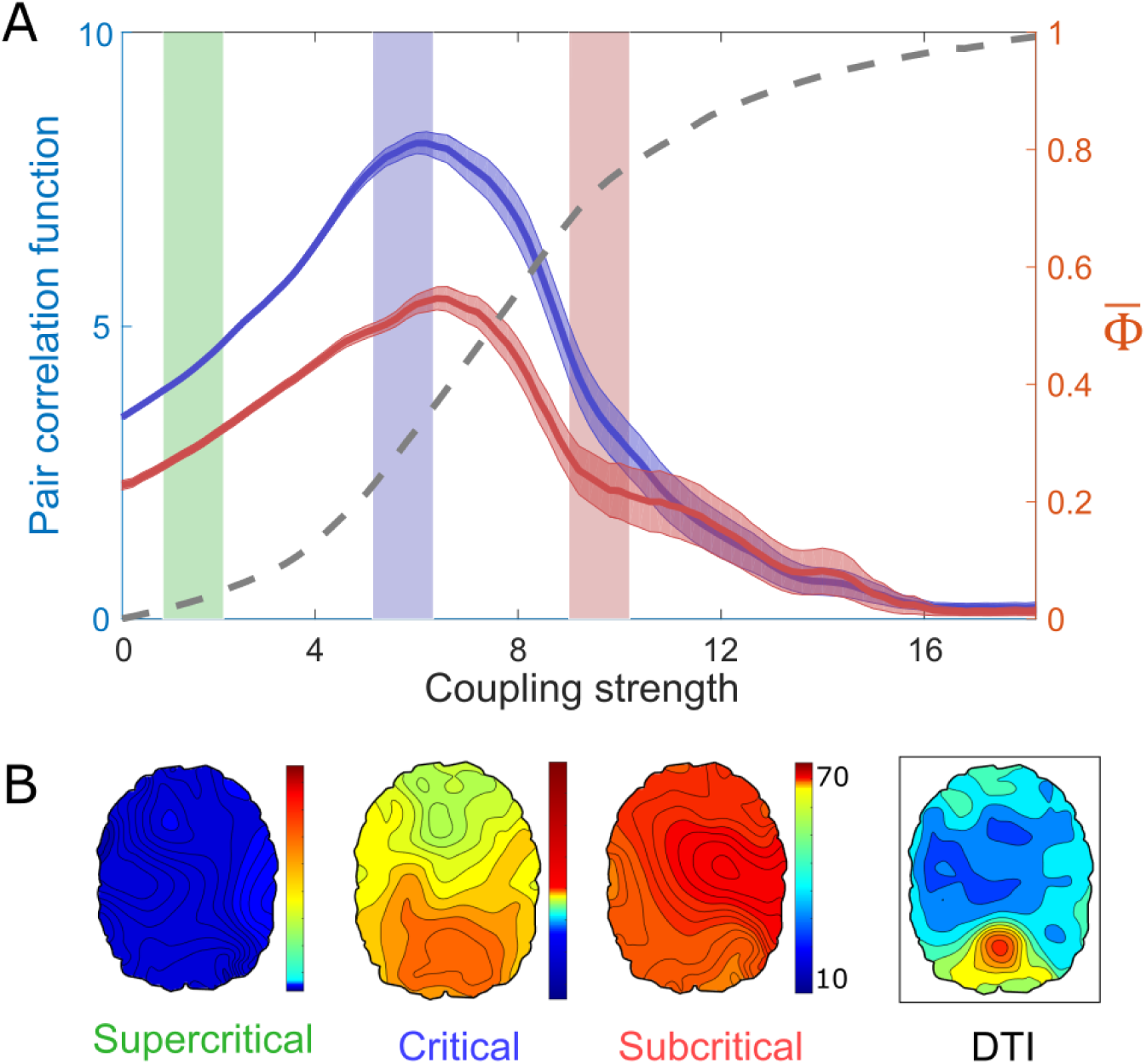
Criticality, 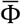, and structured functional connectivity in the brain network. (A) When modulating the coupling strength (a control parameter), the maximum 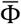 coincides with the maximum criticality, as measured by the pair correlation function (PCF), while the order parameter (dotted line) increases in a monotonic way. (B) Only in the critical state (blue region in Figure 2A), a salient structured functional connectivity, which resembles the structural brain network, emerges in the brain network. Color bar indicates the node degree.

Here, we assumed that the maximum 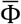 may arise due to the balance between the large variation of functional network configurations and the large constraints of structural network configurations, as a characteristic of a critical state. The disrupted balance in an incoherent or highly synchronized brain network results in a small 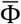. Figure 2B presents the functional brain networks based on the phase lag index (*PLI*). The functional connectivity at the maximum PCF (in the critical state) resembles the structural brain network of DTI, whereas the functional connectivity at the lower PCFs (in the sub-and supercritical states) are relatively homogeneous and not similar with the structural brain network.

### A Network Mechanism of the Maximal 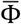 in a Critical State

In a critical state, a network synchronization is balanced by incoherent and synchronous connections through partial phase locking. At a coarse-grained level, the distribution of incoherent and synchronous connections is shaped by the network topology; oscillations at the nodes with dense connections become slower and more synchronous, while oscillations at the nodes with sparse connections are faster and incoherent [25,26]. As a consequence, a coarse-grained functional network resembles a structural network in a critical state [14,54,55].

Figure 3A presents the Spearman correlation coefficients between the node degrees (in the structural brain network) and the *PLI*s of the 78 nodes (in the functional brain network) as coupling strength increases. The Spearman correlation coefficient is at a maximum in the critical state (blue region), while both the super- and subcritical states (green and red regions, respectively) are associated with smaller correlations. The results imply that the constraint of the structural network on a functional network is maximized in the critical state, with higher degree nodes having a larger *PLI*. Figure 3B presents the scatter plots of the *PLIs* versus the node degrees of the 78 nodes in the three states. A large positive correlation coefficient appears only in the critical state (Figure 3B blue, r=0.57, p-value < 0.001). However, the large correlation does not mean that the functional network is static. Figure 3C presents the temporal evolution of the Spearman correlation coefficients between the instantaneous phases of alpha oscillations in the functional brain network and the node degrees in the structural brain network for the three states. The varying correlation coefficients indicate the temporal change of the phase lead-lag relationships among 78 brain regions upon the structural brain network. The large variation of functional networks is one of the characteristics of a critical state and measured by a large PCF. As a result, the large variation of the functional brain network at a small-time scale (∼ seconds) (Figure 3C) under the large constraint from the structural brain network (Figure 3B) in a large time scale (∼ minutes) in the critical state may be the network condition for the maximal 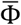 in the brain network.

**Figure 3.**
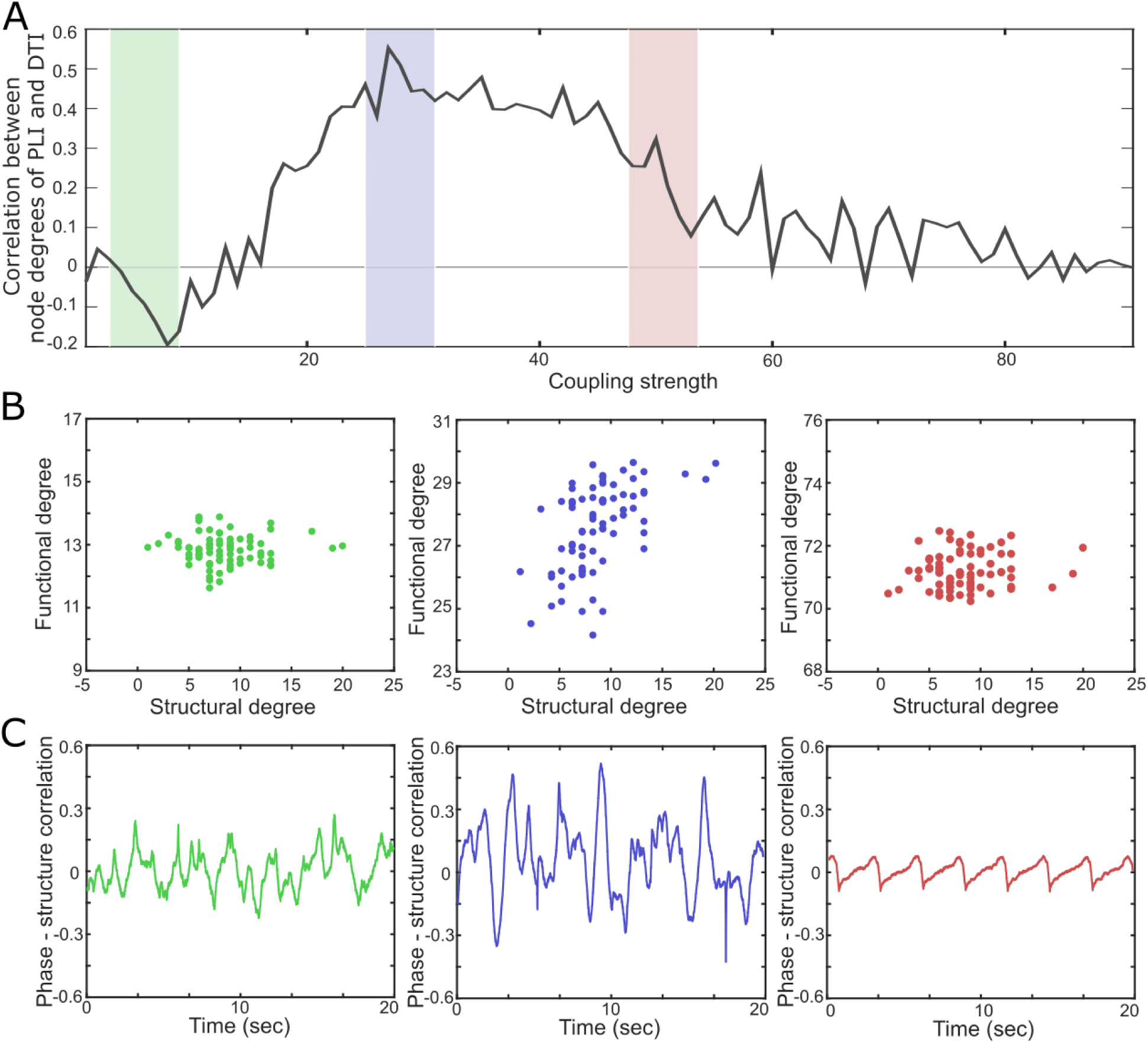
Correlations between the functional and structural brain networks near to and far from a critical state. (A) The Spearman correlation coefficient between the 78 phase lag index (*PLI*) values in the functional brain network and the 78 node degrees in the structural brain network is maximal in the critical state (black line, blue shaded region). (B) The scatter plots (the 78 *PLI* values versus the 78 node degrees) for the supercritical (green), critical (blue), and subcritical (red) states. The large correlation in the critical state implies a large constraint of the structural network on the functional network. (C) The Spearman correlation coefficients between the instantaneous phases of alpha oscillations and the node degrees of the 78 nodes in the structural brain network. The large temporal variation indicates a large repertoire of functional connectivity, which is a characteristic of a critical state.

### EEG study: Correlation between Criticality, 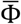, and Human Consciousness

To investigate the relationship between criticality, integrated information, and level of human consciousness, we compared the PCF and 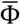 of high-density EEG and the response rate during general anesthesia. The behavioral response rate, which is inversely proportional to the drug concentration, was used as a surrogate for the level of consciousness. In figure 4A, the conscious resting state and the conscious recovery state have a higher PCF and 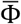 than in the unconscious states. For the continuous EEG data, the change of PCF correlates with the change of 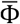. Moreover, both the measures reflect the response rate during the significant state transition. These results are consistent with the model prediction and empirically demonstrate for the first time a direct relationship between PCF, 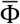, and the level of human consciousness.

**Figure 4.**
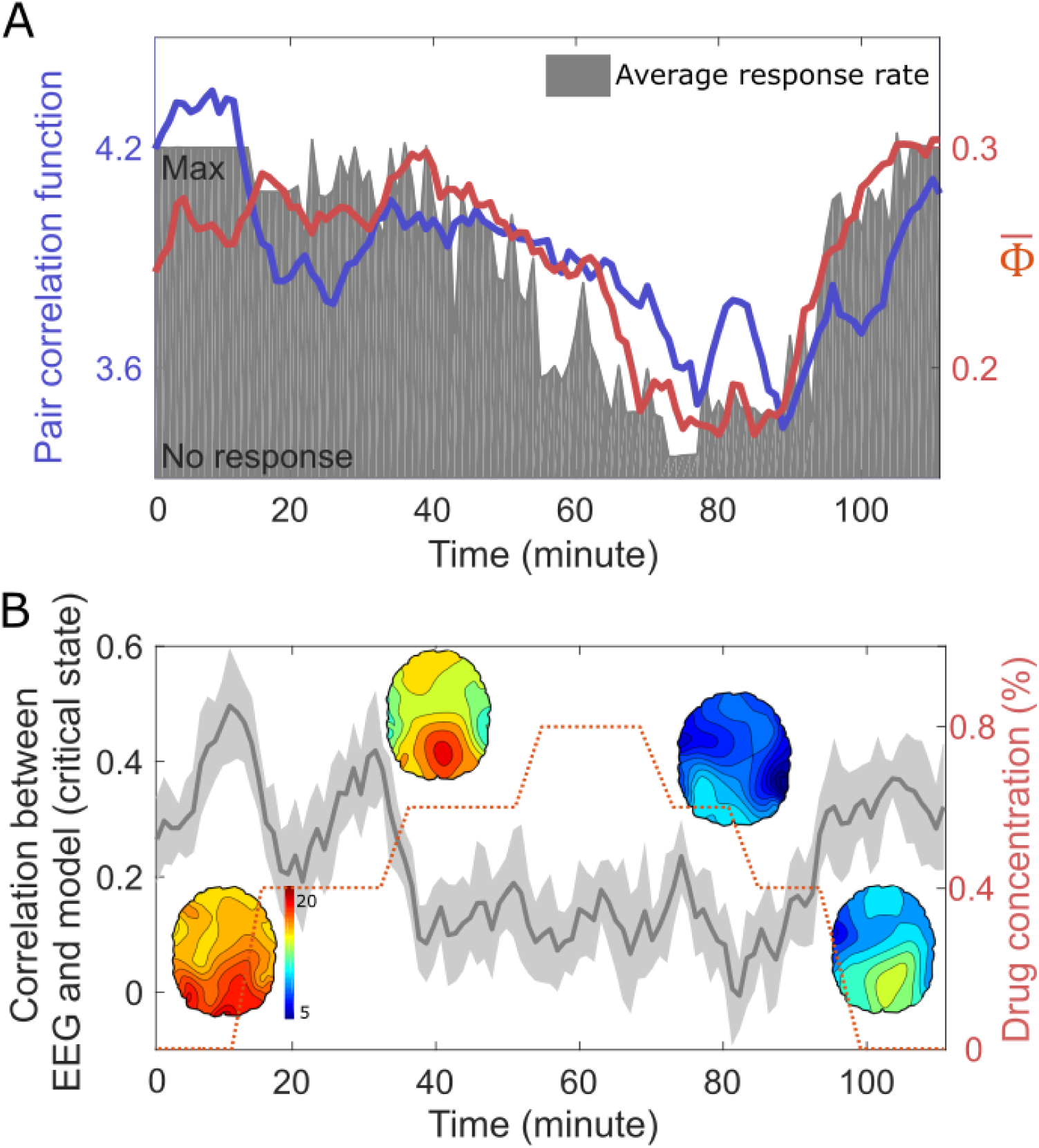
Criticality, integrated information, and level of human consciousness during general anesthesia. (A) The PCF and 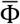 of 64-channel EEG correlate with the response rate (grey area), which was modulated with increasing anesthetic concentrations. (B) The Spearman correlation coefficients between the *PLI* networks of the EEG and the simulated EEG based on the anatomical brain network and critical state. The conscious states (baseline, induction, and recovery) show larger correlations, which implies a stronger constraint of the structural brain network on the EEG in conscious states. As a result, the balance between the large constraint of the structural brain network and the large repertoire (i.e., large PCF) of the functional brain network may be the network basis of the large 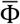 of EEG in the conscious brains.

The model study also predicted a large correlation of the structural brain network with the functional brain network in consciousness, which corresponds to the functional brain network in a critical state. However, since we recorded only the scalp EEG, we were not able to test the correlation between functional (*PLI*) and structural (node degree) brain networks. Instead, we compared the *PLI* networks of the EEG and a simulated EEG. For the simulation of EEG in the conscious state, we first simulated the source signals in the structural brain network in a critical state (the same simulation in Figure 2 and 3) and then projected the source signals onto the scalp (see the Method in details). If the model prediction is correct, both the *PLI* networks should be largely correlated.

Indeed, we found high Spearman correlation coefficients between the *PLI* network of the EEG in the conscious states and the *PLI* network of the simulated EEG in a critical state. This correlation decreases along with the reduced response rate in the unconscious states (Figure 4B). The high correlation coefficients in conscious states indicate a strong constraint of the structural brain network on the EEG.

## Discussion

### Summary of the findings

In this paper, we performed a computational modeling and EEG data analysis to study the relationship between integrated information, criticality, and level of human consciousness. In the modeling study, we simulated the brain network activities near and far from a critical state and quantified the criticality and integrated information with PCF and 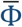. We found that when the brain network reaches a maximal PCF, the 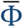 is also maximized under the largest constraint of the structural brain network (that is, with the largest correlation between the functional and structural brain networks). To verify the modeling results, we gradually modulated the level of consciousness in humans with a general anesthetic and compared the PCF and 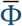 of high-density EEG with the behavioral response rate. We found that, in the conscious resting states, the subjects have the highest PCF and 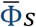 when the behavioral response rates are highest. We also showed that the functional brain network of EEGs in conscious states largely correlates with the functional brain network of the simulated EEGs, which were modeled to reflect the underlying structural brain network in a critical state. From both the modeling study and EEG analysis, we propose that the balance between the large variation of functional brain network (as measured as the large PCF) and the large constraint from the structural brain network in a critical state is a necessary condition to generate the high 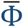 of conscious states.

### Criticality and maximal 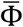

IIT’s composition principle states that any subset of elements within a system can be a mechanism of the system if the intrinsic cause-effect power is irreducible [1,3]. Furthermore, the set of all mechanisms and their constraints within the system comprises a system’s cause-effect structure. Algorithmically, the intrinsic cause-effect structure can be identified by the effective information among partitions (*φ*), with the integrated information of a system (Φ) defined by the information transmission of the minimal information bipartitions. However, in the application to our brain network model, since we did not give any cause-effect structure initially, a non-zero 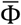 may emerge through only the interaction of 78 nodes in the brain network. This raises the following questions. How does a non-zero 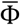 emerge spontaneously in the brain network? What is the network basis that facilitates the emergence of a cause-effect structure? In this study, we propose that the phase lead-lag relationship that is shaped by the structural brain network may create conditions that facilitate the emergence of a cause-effect structure.

Previous studies have examined how brain network topology modulates the frequencies and phases of local node dynamics, subsequently shaping a pattern of global information flow and functional connectivity. A mathematical relationship between the node degree and the phase of node dynamics was identified analytically [25,26], which enables us to estimate analytically the phase of a node (in a functional network) with only its node degree and local connectivity (in a structural network). This was tested with diverse brain networks (human, monkey, and mouse), which demonstrated that when considering long-term and spatially coarse-grained brain activities (> minutes) the global phase lead-lag relationship and information flow pattern (measured by transfer entropy and Granger causality) were predictable based only on the structural brain network topologies of the three species. In particular, the difference between the “hub node” and “peripheral node” becomes most salient at a critical state. In other words, the structural complexity of overall coupled node dynamics is maximized at a critical state. The differences between frequency and phase among coupled node dynamics naturally give rise to information flow in the brain network, consequently dividing the nodes in the brain network into ‘senders’ and ‘receivers’ of information flow. However, in sub- and supercritical states, there is no information flow between node dynamics because of a highly-synchronized state (i.e., no difference that would allow for information flow) and incoherent state (i.e., no interaction that would allow for information flow). Therefore, a cause-effect structure may emerge between these two extreme states and be maximized at a balanced state. Extrapolating from results based on the phase lead-lag relationship, emerging a cause-effect structure without external stimuli is likely not random but rather is organized by a mathematical relationship between network structure and dynamics in a critical state [25,26,56,57].

### A large variation under a large constraint

In our modeling study and EEG analysis, we found that a large variation of the functional brain network occurs in spite of a strong constraint from the structural brain network. Notably, the large variation and the strong constraint were observed at different time scales. When we investigated the correlation coefficient between the instantaneous phases and the node degrees of the 78 nodes, the correlation coefficient at each time point varied widely at a short time scale (∼ seconds) (Figure 3C). By contrast, the averaged instantaneous phases and the node degrees at a longer time scale (∼ minutes) have a large positive correlation coefficient (r=0.57, Figure 3B), which implied a bias toward positive values in the short time scale variation. Our previous EEG study demonstrated that the correlation coefficient between the EEG network and the structural brain network was pronounced when calculated with large temporal windows (> 60 seconds) but diminished when calculated with small windows (< 5 seconds). Interestingly, the scale-dependency appeared only in the baseline conscious states, not in the altered states of consciousness such as anesthetized state, minimally conscious state, and unresponsive wakefulness syndrome [23]. The conscious brain can be characterized by diverse repertoires of global functional connectivity on a short time scale. However, when combining all repertoires of small windows in a large window, the different functional connectivity patterns across small windows may be averaged out and only the common functional connectivity patterns that are constrained by the structural brain network remain. Such an averaged functional connectivity in a large window can be simulated by the mean-field method upon a structural brain network. Nevertheless, how the large variance and the large constraint contribute to the large integrated information in conscious states remains elusive. Further studies are required to understand the mechanistic association between the functional variance, structural constraint, and Φ in a network.

### Deviations from criticality and 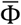

When brain dynamics deviate from a critical state, the capacity to shape global node dynamics into a structure that resembles the network structure is lost. In a supercritical state, there is no functional interaction between nodes. In a subcritical state, node dynamics cannot be structured due to strong functional interactions between nodes that eliminate functional heterogeneity. As a result, the 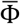 of a brain not in a critical state—regardless of whether the deviation is toward subcriticality or supercriticality—will decrease in proportion to the distance from the critical state.

Many empirical studies support the criticality hypothesis in consciousness by comparing the dynamics of various conscious states with unconscious states (such as sleep, anesthesia, seizure, and coma). They have commonly shown that unconsciousness is associated with a deviation from criticality, which is quantified with various methods (power law, susceptibility, pair correlation function, and correlation between the functional and structural brain networks) [15,22,23,40,58–68]. However, no one has yet associated criticality, human consciousness, and Φ because of the explosive computational demands required for Φ calculations. In a previous study, we introduced a method that statistically estimates the increase/decrease of Φ for a continuous EEG, termed 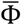, and showed the applicability to characterizing levels of consciousness with high-density EEG and also enabled us to study a relationship with network criticality. Our current results show that the empirical and computational model studies support the hypothesis that the characteristic network properties in a critical state naturally give rise to a structured, asymmetric phase lead-lag relationship among oscillators and, in turn, maximize Φ in the conscious brain.

### Limitations

This study has several limitations. First, 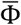 does not measure the precise Φ. Instead, we estimated the relative change of Φ, focusing on the increase/decrease along with the change in criticality in the network model and based on empirical EEG during anesthetic-induced unconsciousness. Even so, this relative measure is sufficient for the purpose of this study, which is to examine the relationship between Φ, criticality, and levels of consciousness. And considering the significant differences among the six versions of Φ in the recent comparison [69], testing our conclusion with other versions of Φ is required. Second, in this study, we derived a relationship between asymmetric phase lead-lag relationships and 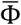, but this relationship does not explain how the phase lead-lag relationship generates a cause-effect relationship defined by information partition for Φ. It may require an analytic study to identify conditions to link phase lead-lag and Φ. Third, to find an association between Φ, criticality, and consciousness, we used the subjects’ response rate during exposure to a general anesthetic as a surrogate for the level of consciousness. However, it is well known that unresponsiveness does not necessarily correlate with lack of consciousness. Although previously studied in the context of vegetative and minimally conscious states, our research team has recently identified covert consciousness during propofol sedation in which overt motoric responses were absent but brain network responses suggestive of volition were present [70]. Future studies might incorporate EEG data, behavioral responsiveness, and neuroimaging protocols to determine covert consciousness in order to more precisely identify the relationship between critical dynamics, consciousness, and Φ. Finally, the structural brain network used in the modeling study includes only the cortex. Including subcortical networks such as thalamocortical and hippocampocortical connections, etc. could improve the modeling performance for complex state transitions during general anesthesia.

### Conclusion

We demonstrated for the first time an explicit relationship between criticality, integrated information, and human consciousness with computational modeling and EEG analysis. We propose that network criticality, that is, a balanced state between large variation of functional network configurations and strong constraint of structural network configurations, is a network condition for integrated information. Understanding this relationship may open a new way to study diverse states of consciousness situated near to and far from a critical state in terms of integrated information. It may also provide a theoretical foundation for controlling the level of consciousness and integrated information by modulating criticality at a network level.

## Acknowledgments

This work is supported by grant No. R01 GM098578 (PIs: GM and UL) from the National Institutes Health.

