## Supplemental text, table1, table2 for "Criticality as a determinant of integrated information Φ in human brain networks"

**1. Experimental procedure of the study of sevoflurane-induced unconsciousness in humans**

The EEG data was recorded from seven healthy volunteers (4 males, 20–23 years of age) who gave their written informed consent participated in the study. Participants were American Society of Anesthesiologists class 1 physical status, body mass index less than 30, with Mallampati 1 or 2 airway classifications and participants who were pregnant, or with a history of obstructive sleep apnea, gastroesophageal reflux, cardiac conduction abnormalities, asthma, epilepsy, history of problems with anesthesia, family history of problems with anesthesia, history of drug use, and any neurologic or psychiatric history were excluded from the study. Participants kept their eyes closed during the experiment. Sevoflurane was administered by a secured face mask at an initial concentration of 0.4% in high-flow oxygen (8 L/min). After 15-min equilibration of each concentration, the EEG data were recorded during 10-min at the target concentration. The concentration of sevoflurane was increased by levels of 0.2% until the participants lose their responsiveness(LOR). After 10-min period of LOR, the reverse protocol was proceed until the responsiveness is recovered. The participants were instructed to squeeze objects in each hand every 30 s for assessing responsiveness. 64-channels sensor cap from Electrical Geodesics, Inc. was used to record EEG with a sampling frequency 500Hz. The impedance of channels was reduced to below 50 KΩ before data acquisition.

**2. Supplementary Tables**

**Supplementary Table 1.** Statistical tests of Pair Correlation Function(PCF) and integrated information(Φ) among supercritical, critical, and subcritical states in model

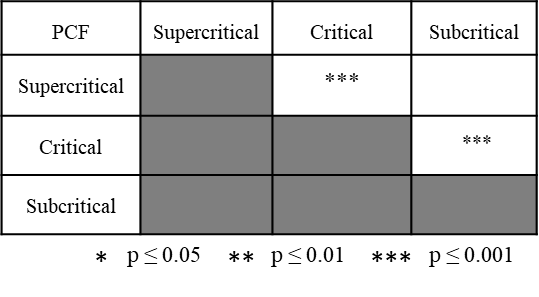

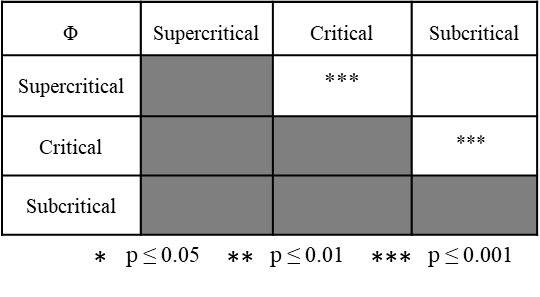

**Supplementary Table 2.** Statistical tests of Pair Correlation Function(PCF) and integrated information(Φ) among baseline, induction, anesthesia, and recovery states in experiment

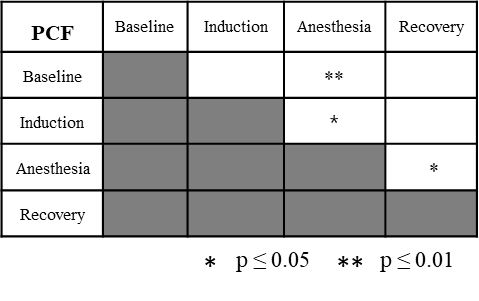

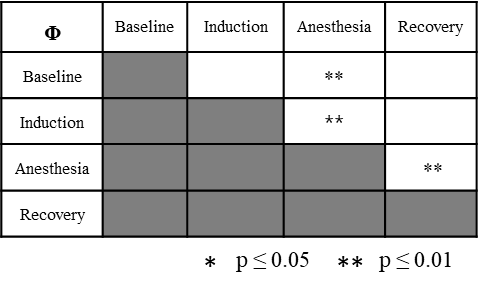

The baseline, induction, anesthesia, and recovery states are the average of 10-min with 0%, 0.4%, 0.8%, 0% drug concentrations, respectively.
